# Laue-DIALS: open-source software for polychromatic X-ray diffraction data

**DOI:** 10.1101/2024.07.23.604358

**Authors:** Rick A. Hewitt, Kevin M. Dalton, Derek Mendez, Harrison K. Wang, Margaret A. Klureza, Dennis E. Brookner, Jack B. Greisman, David McDonagh, Vukica Šrajer, Nicholas K. Sauter, Aaron S. Brewster, Doeke R. Hekstra

**Affiliations:** Department of Molecular and Cellular Biology, Harvard University, Cambridge, MA 02138; Linac Coherent Light Source, SLAC National Accelerator Laboratory, Menlo Park, 94025, CA, USA; New York University, New York, NY 10012; SLAC National Accelerator Laboratory, Menlo Park, CA 94025; Graduate Program in Biophysics, Harvard University, Boston, MA 02115; Department of Chemistry and Chemical Biology, Harvard University, Cambridge, MA 02138; Science and Technology Facilities Council, Rutherford Appleton Laboratory, Didcot, OX11 0FA, United Kingdom; BioCARS, Center for Advanced Radiation Sources, The University of Chicago, Chicago, Illinois 60637, USA; Molecular Biophysics and Integrated Bioimaging Division, Lawrence Berkeley National Laboratory, Berkeley, CA 94720 USA; School of Engineering and Applied Sciences, Harvard University, Allston, MA 02134

## Abstract

Most X-ray sources are inherently polychromatic. Polychromatic (“pink”) X-rays provide an efficient way to conduct diffraction experiments as many more photons can be used and large regions of reciprocal space can be probed without sample rotation during exposure—ideal conditions for time-resolved applications. Analysis of such data is complicated, however, causing most X-ray facilities to discard *>*99% of X-ray photons to obtain monochromatic data. Key challenges in analyzing polychromatic diffraction data include lattice searching, indexing and wavelength assignment, correction of measured intensities for wavelength-dependent effects, and deconvolution of harmonics. We recently described an algorithm, Careless, that can perform harmonic deconvolution and correct measured intensities for variation in wavelength when presented with integrated diffraction intensities and assigned wavelengths. Here, we present Laue-DIALS, an open-source software pipeline that indexes and integrates polychromatic diffraction data. Laue-DIALS is based on the dxtbx toolbox, which supports the DIALS software commonly used to process monochromatic data. As such, Laue-DIALS provides many of the same advantages: an open-source, modular, and extensible architecture, providing a robust basis for future development. We present benchmark results showing that Laue-DIALS, together with Careless, provides a suitable approach to the analysis of polychromatic diffraction data, including for time-resolved applications.

## I. INTRODUCTION

Most X-ray generation mechanisms inherently yield polychromatic (“pink”) X rays, with the energies of incident photons often varying by several percent (Table I). Historically, analysis of such data has been complicated, leading many X-ray facilities to discard *>*99% of X-ray photons to obtain monochromatic data. While this simplifies analysis, pink X-ray diffraction can be a necessary stage in the deployment of new X-ray generation technologies that do not yet provide sufficient flux for successful monochromatic data collection. More fundamentally, polychromatic X-rays cover significant regions of reciprocal space while recording full rather than partial diffraction intensities without the need for sample rotation during exposure, in contrast with monochromatic exposures. This opens up applications in which samples cannot be rotated during exposure, for instance with short X-ray pulses, while also increasing the number of incident photons. For these reasons, pulsed polychromatic X-rays are ideally suited to the study of the dynamics of biological macromolecules–a major frontier in structural biology.

**TABLE I.**
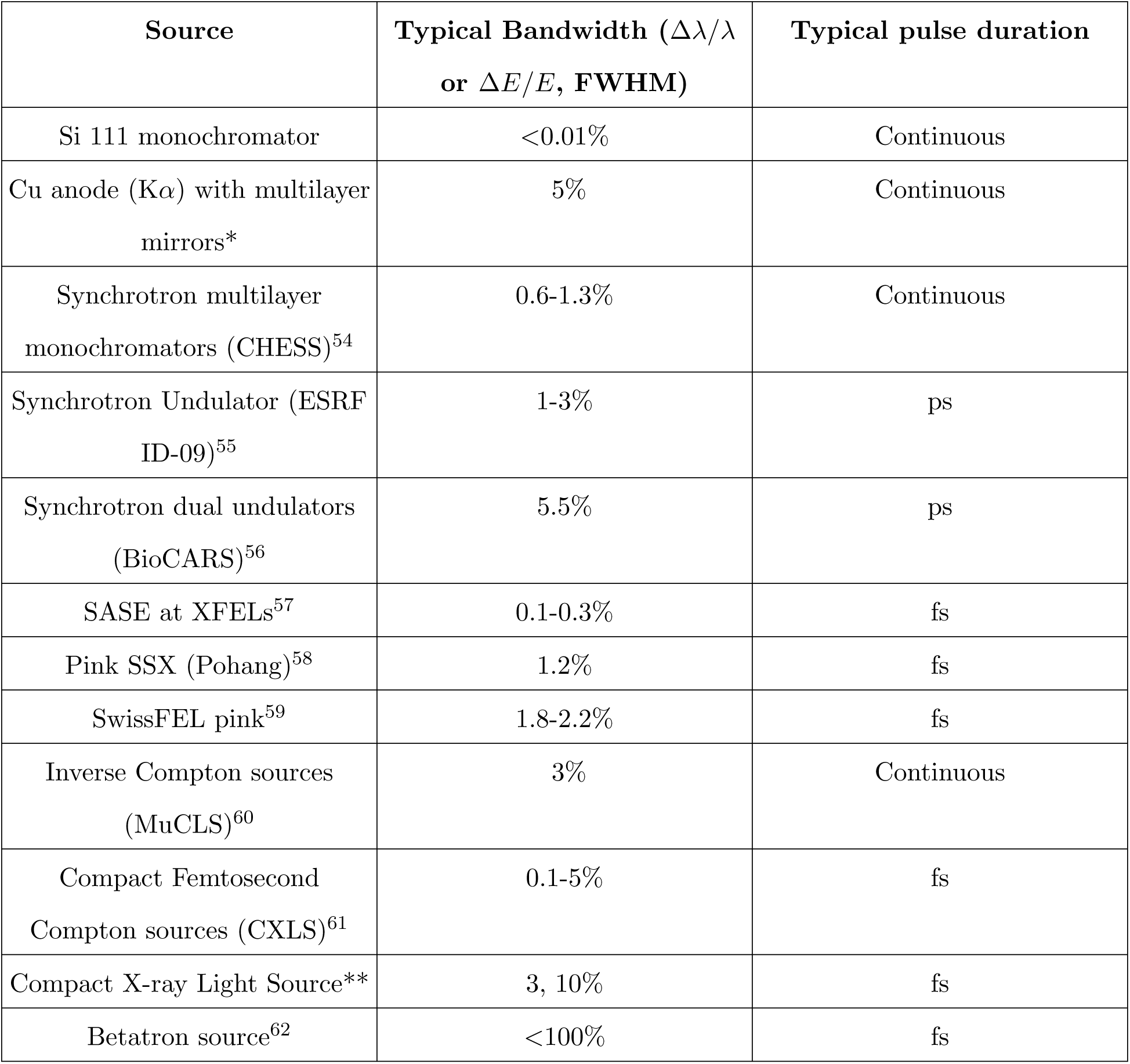
Example instrument X-ray bandwidths. * Based on personal communication with Dr. J. Graf, Oncoatec (Bruker). ** Based on personal communication with Dr. W.G. Graves, Arizona State University.

| Source | Typical Bandwidth ($\Delta\lambda/\lambda$<br>or $\Delta E/E$, FWHM) | Typical pulse duration |
| --- | --- | --- |
| Si 111 monochromator | <0.01% | Continuous |
| Cu anode ($K\alpha$) with multilayer<br>mirrors* | 5% | Continuous |
| Synchrotron multilayer<br>monochromators (CHESS)<sup>54</sup> | 0.6-1.3% | Continuous |
| Synchrotron Undulator (ESRF<br>ID-09)<sup>55</sup> | 1-3% | ps |
| Synchrotron dual undulators<br>(BioCARS)<sup>56</sup> | 5.5% | ps |
| SASE at XFELs<sup>57</sup> | 0.1-0.3% | fs |
| Pink SSX (Pohang)<sup>58</sup> | 1.2% | fs |
| SwissFEL pink<sup>59</sup> | 1.8-2.2% | fs |
| Inverse Compton sources<br>(MuCLS)<sup>60</sup> | 3% | Continuous |
| Compact Femtosecond<br>Compton sources (CXLS)<sup>61</sup> | 0.1-5% | fs |
| Compact X-ray Light Source** | 3, 10% | fs |
| Betatron source<sup>62</sup> | <100% | fs |

Such diffraction images collected without sample rotation are known as stills. The analysis of monochromatic stills is particularly fraught by the partiality problem. When samples are rotated (Figure 1a), the complete intensity of reflections can be collected. Without rotation, only a small part of the intensity of reflections is observed (Figure 1b). In addition, many fewer reflections are typically observed. Several analytical approaches have been proposed to alleviate the partiality problem, but none are accurate enough to remove the requirement for large numbers of diffraction patterns to achieve accurate structure factor amplitudes.

**FIG. 1.**
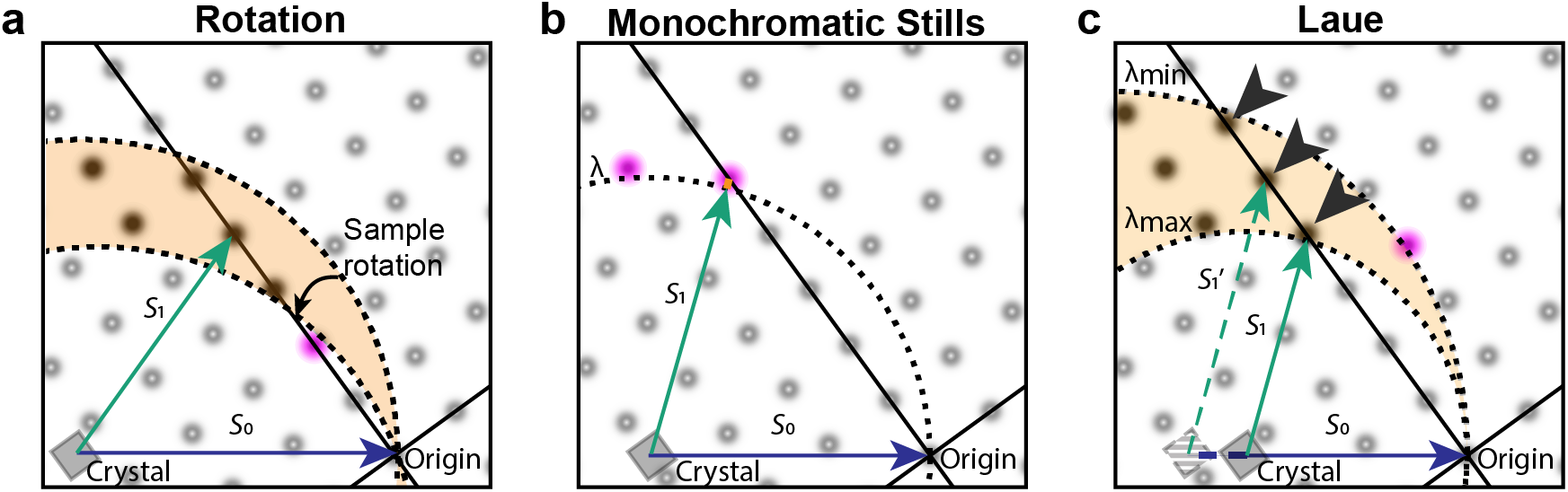
Pink and monochromatic data collection. **A.** Ewald diagram for an exposure during a conventional monochromatic exposure during sample rotation (not to scale). Rotation of the sample corresponds to rotation of the reciprocal crystal lattice around its origin (black dot, right corner), sweeping out a corresponding section (orange) of the reflections (reciprocal lattice points). Reciprocal lattice points shown as fuzzy dots; observable reflections, which pass through the Ewald sphere, are shown as darker dots. Partially observed reflections are shown in pink. **B.** For monochromatic stills, few reflections are observable, and for those that are observed, intensities are only partially observed. **C.** The Laue method collects polychromatic stills using a range of wavelengths from *λ*_min_ to *λ*_max_. As a result, more reflections are observed and mostly at their full intensity. Incident beam vectors (*S*_0_) shown in blue; scattered beam vectors (*S*_1_) in green. Variation in the length of *S*_0_ and *S*_1_ vectors in panel C indicates that these correspond to different wavelengths. Harmonics are indicated with arrowheads and lie on “central rays” which pass through the origin of the reciprocal lattice. Partials are rare and occur predominantly at low resolution—close to the origin of the reciprocal lattice.

The collection of stills by exposure of crystals to polychromatic X-rays is known as the Laue method^1^. As illustrated in Figure 1c, the Laue method largely eliminates the partiality problem: most reflections will be observed in full, with only the need to correct for the variation in diffracted intensity with wavelength. Many applications benefit, including synchrotron serial and time-resolved crystallography, and experiments at X-ray Free Electron Laser (XFEL) facilities. The latter experiments are usually based on the diffract-before-destroy principle as individual femtosecond XFEL exposures often exceed the radiation damage dose limit by orders of magnitude^2,3^. Such XFEL experiments have enabled determination of structures from sub-micron-sized crystals^4^ and of damage-free structures of metalloproteins^5^, as well as a wealth of time-resolved studies^6–11^.

The Laue method is, in principle, an ideal approach for any crystallographic technique requiring the collection of stills. Despite the attractive properties of the Laue method, however, it has found limited adoption outside of synchrotron time-resolved crystallography studies. There are a few reasons for this. First, the appearance of Laue diffraction patterns is highly sensitive to crystal mosaicity (Figure 2, top row), complicating lattice search and integration of the intensity per spot. Second, the options for processing macromolecular Laue diffraction patterns are limited and do not readily permit incorporation of new algorithms for performing different aspects of the data reduction pipeline, such as new indexing and integration algorithms. For instance, Precognition (Renz Research, Inc.) is proprietary, closed-source software, while the Daresbury Laue suite^12^ does not readily compile on modern hardware or interface with scientific Python.

**FIG. 2.**
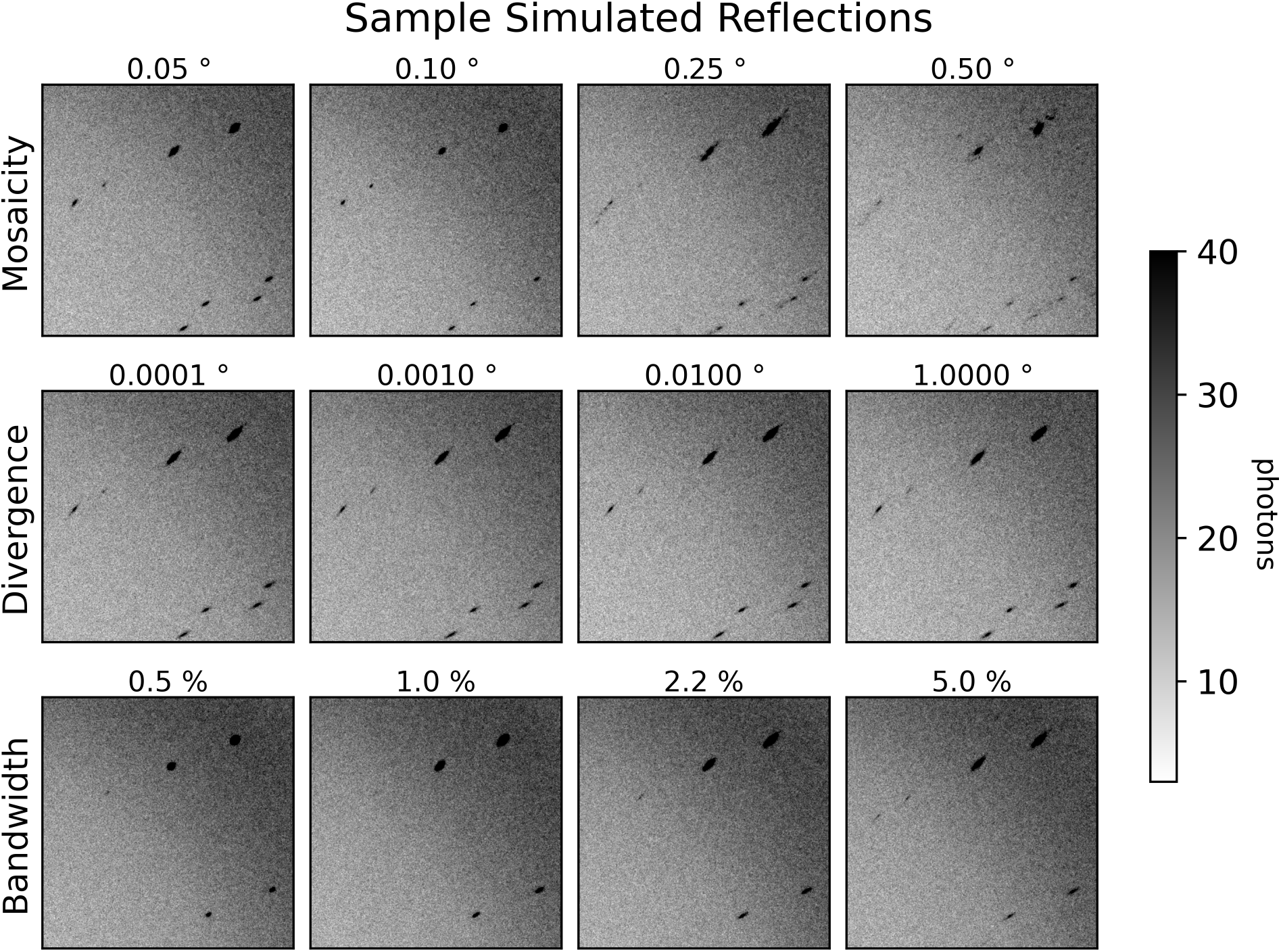
Dependence of spot profiles on simulated crystal parameters. Close-ups of synthetic diffraction patterns for dihydrofolate reductase, generated by nanoBragg for different levels of crystal mosaicity, beam divergence, and beam spectral bandwidth. An input BioCARS spectrum of 5.5% bandwidth at full-width half-maximum is used for all simulations except those that vary bandwidth, which use a simulated Gaussian spectrum with the same center and varying bandwidths. The reference data set here is labeled with 0.1*^◦^* mosaicity, and uses the input BioCARS spectrum with 0.1*^◦^* beam divergence. Each other data set differs from that data set by only the associated label.

Figure 1 also illustrates several of the difficulties in interpreting Laue diffraction patterns. In the monochromatic case, each reflection can be mapped fairly accurately to the reciprocal lattice, as the lengths of the *S*_0_ and *S*_1_ vectors are known (the inverse of the wavelength; panels A and B). For Laue diffraction (Figure 1c), different reflections lead to diffraction at the same time but for different X-ray wavelengths (evident as variation in the length of the *S*_0_ and *S*_1_ vectors). As a consequence, the mapping is not unique, as the wavelength is not observed (except in neutron time-of-flight Laue diffraction). This complicates crystal lattice determination, assignment of Miller indices, and inference of the wavelength of each observed diffraction spot. In addition, observed intensities need to be corrected (“normalized”) for dependence of incident flux, absorption, and diffraction on the wavelengths of the X rays for which the diffraction condition is met (along with corrections for other effects such as exposed crystal volume and radiation damage). Finally, reflections with the same direction of the *S*_1_ vector (marked by arrow heads in Figure 1c) will be coincident on the detector—this occurs for reflections that lie on the same central ray (lines passing through the origin of the reciprocal lattice). Such diffraction spots with contributions from multiple reciprocal lattice points are known as harmonics.

Here, we introduce Laue-DIALS: an open-source, extensible, Python-based platform for the reduction of Laue data to integrated intensities. At present, the package enables the processing of fixed-target pseudo-rotation series data—conventional single-crystal Laue data collected as a set of stills at a series of evenly-spaced angles. In doing so, Laue-DIALS addresses lattice determination, assignment of indices and peak wavelengths per reflection, geometry refinement, and integration. Importantly, Laue-DIALS provides a general platform for further development and deployment of novel algorithms, including those based on machine learning—a natural pairing given the spectral complexity of Laue diffraction patterns. Laue-DIALS naturally interfaces with the program Careless, an open-source, Python-based software^13^. Careless performs simultaneous scaling and merging of X-ray diffraction data using forward probabilistic modeling of data formation and addresses wavelength normalization and the deconvolution of harmonic observations.

Laue-DIALS builds on DIALS (Diffraction Integration for Advanced Light Sources)^14^. DIALS is an open source toolkit for working with diffraction data. Its components include a library (dxtbx^15^) that is able to read images from the vast majority of known X-ray sources, and to represent beamline geometry using programmatic, parameterized models (such as beam, detector, crystal, goniometer, and scan). DIALS also includes a set of routines for searching images for bright Bragg signal (spot finding), determining the crystal orientation and unit cell parameters from the spot locations (indexing), refinement of crystallographic models^16^, prediction of spot centroid locations, and integration of pixels to create summed measurements of Bragg reflections. DIALS was developed from the ground up to be flexible, extensible, and reusable, with a toolkit approach to its libraries to support modification and addition^17^. Critically, it was based on algorithms developed and kindly published by crystallographers (such as XDS^18^ and MOSFLM^19^), and as such is robust in use cases ranging from standard rotation crystallography, to serial crystallography^20^, to micro-electron diffraction^21^. DIALS is used at many beamlines for routine data processing, both in manual and automated pipelines to process hundreds of thousands of datasets per year.

DIALS originated with the monochromatic diffraction experiment in mind, but direct support of polychromatic sources (including X-ray and neutron sources) is in development. In the meantime this work uses DIALS for reading in Laue data, spot finding, generating an initial indexing solution, and refinement of experimental geometry. Afterwards, new code described here is used for Laue specific steps, such as wavelength assignment and normalization.

## II. THE LAUE-DIALS DATA REDUCTION WORKFLOW

We illustrate the typical workflow for processing Laue diffraction data using Laue-DIALS in Figure 3.

**FIG. 3.**
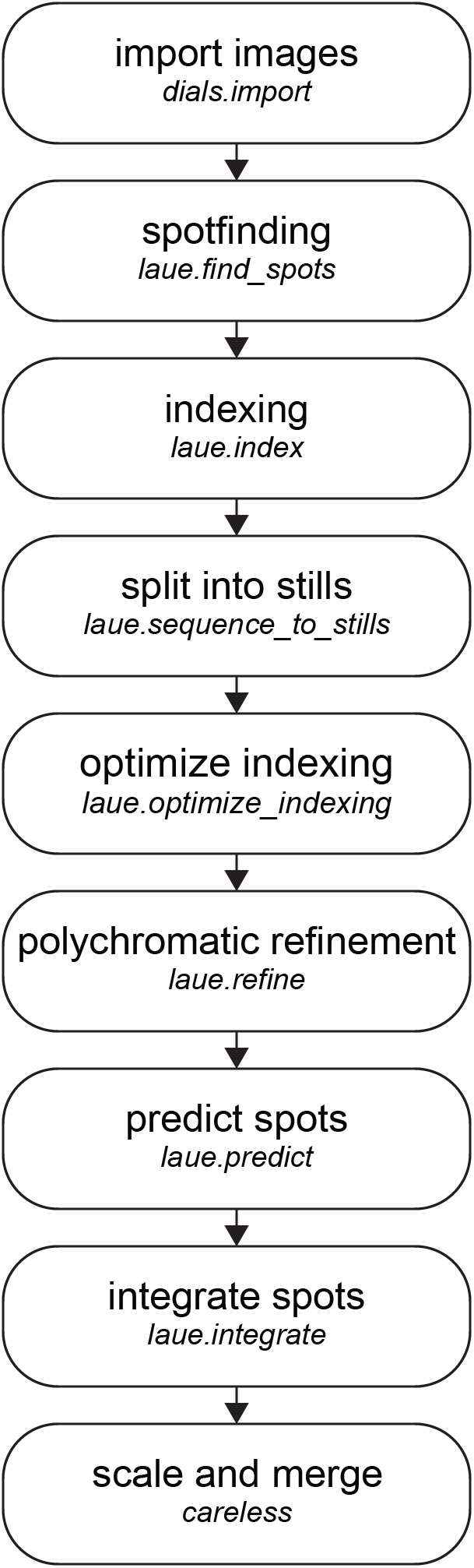
The standard Laue-DIALS data reduction pipeline. Raw image data are imported into a DIALS file format with dials.import, with spot finding and monochromatic indexing to follow. The data are then split into still images, and laue.optimize indexing performs wavelength assignment for each reflection while jointly refining the crystal orientation per image. Geometric refinement produces a final experimental model which is used to predict spot centroids for all reflections likely to contribute to each image. Integrating the reflections then provides a set of integrated intensities and associated metadata usable in Careless for scaling and merging.

### A. Importing data

The current Laue-DIALS analysis pipeline begins with application of monochromatic algorithms present in DIALS to generate an initial estimate of the experimental geometry. Experimental data are imported using dials.import, storing the initial parameter values of the beam, detector, image set, and goniometer models in a so-called experiment (.expt) file. At the software level, these parameterized models take the form of objects as described by Waterman et al.^22^ At this stage a pixel mask can be provided that will be applied automatically at all subsequent steps, or such a mask can be provided at individual subsequent steps to mask panel gaps, dead panels, beam stop shadows, etc. We advise users to construct masks using the DIALS image viewer to suit their use case.

### B. Spot finding, indexing, and initial refinement

The next Laue-DIALS commands are thin wrappers of their DIALS counterparts. These commands override certain parameters with appropriate defaults for polychromatic data. The overridden parameters for each of these commands are enumerated in Table II. For spot finding, we use a 2-D spot finder in DIALS to locate reflection centroids on each image individually. Although we do not override spot finding gain or masking, these parameters are critical for success, and we recommend that the user use the DIALS image viewer to assess if spots have been identified appropriately, leaving neither too many clear reflections unidentified nor marking pixel noise as reflections.

**TABLE II.**
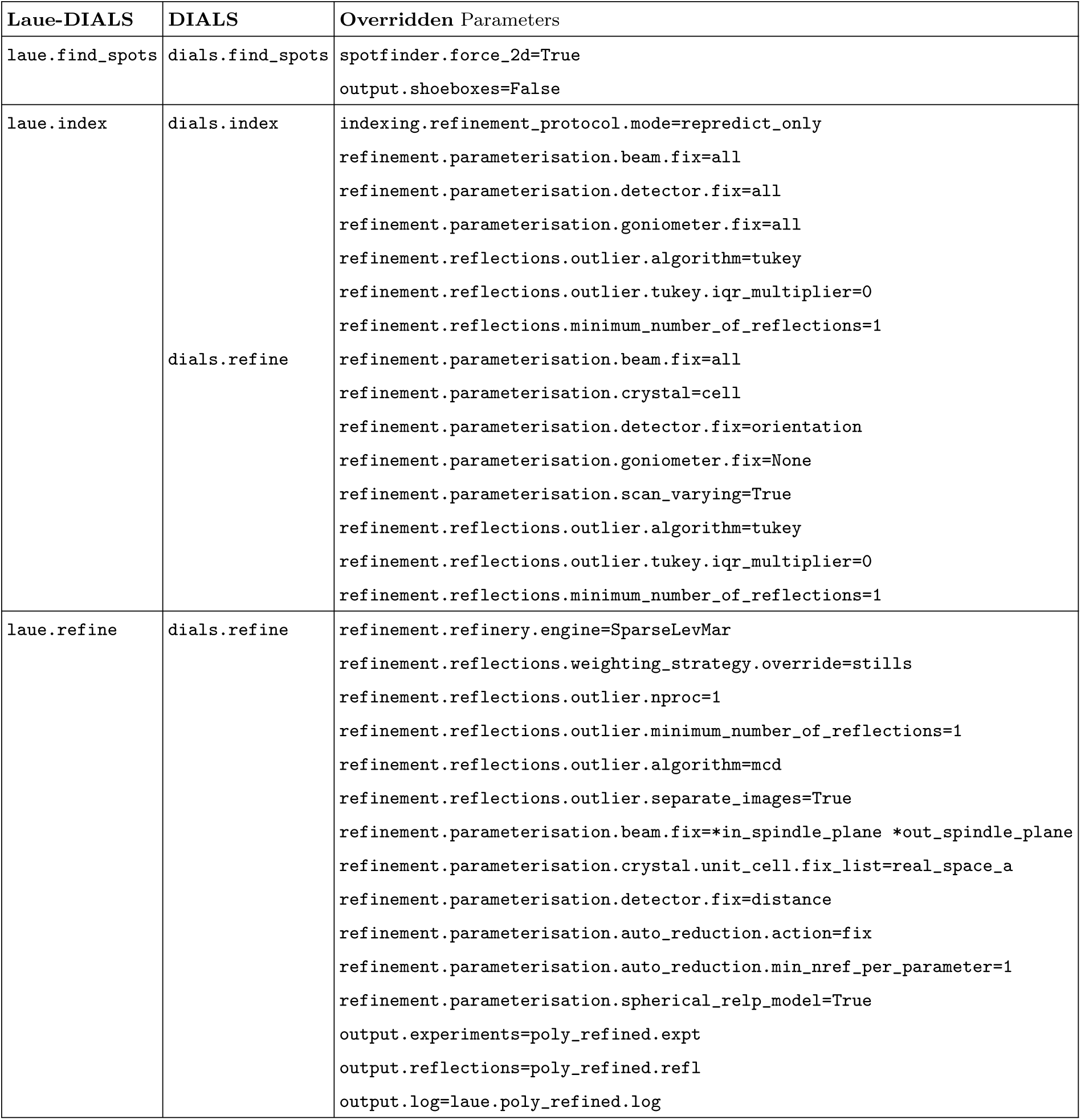
Overridden parameters for Laue-DIALS wrappers. All DIALS parameters that are overridden by Laue-DIALS for wrapper programs. Valid as of DIALS v3.17.0 and Laue-DIALS v0.4.

To index the reflections found by spot finding, we apply the FFT3D algorithm implemented in DIALS^14^ at a nominal wavelength matching either the peak spectral intensity or midpoint of the beam spectrum. All available images are used for indexing. This algorithm determines the unit cell and orientation for the crystal, assigns Miller indices to a subset of the reflections (typically 10-25%), and assigns these reflections to the nominal wavelength. At this point, this subset may correspond to a true wavelength that differs from the specified wavelength, causing a multiplicative shift in the inferred unit cell dimensions––this can be accounted for later (see below). If the unit cell and space group of the crystal are known, those can be provided at this indexing step to improve the initial estimate. By default, we override the input parameters, fixing all beam, detector, and goniometer parameters, allowing only for the crystal parameters to be estimated. Tukey outlier rejection^23^ is used to disregard reflections that do not approximately fit an indexed experimental model during iterations of indexing. An optional iteration of geometric refinement can also be applied, which takes one of two forms: scan-static or scan-varying. For experiments where geometric parameters are expected to vary smoothly across the dataset, a round of scan-varying refinement can improve the accuracy of the solution by allowing variation of geometric parameters across images in a given dataset. For experiments where variation of parameters is unknown or expected not to vary, the scan-static option allows for a single universal experimental geometry to be applied equally to all images. This initial geometric refinement in Laue-DIALS still fixes all aspects of the beam, orientation of the detector, and unit cell of the indexed crystal (but not orientation), but allows for free variation of the goniometer parameters. Tukey outlier rejection is also applied in this stage. Any of the parameters listed here can be overridden by the user, including those that Laue-DIALS overrides itself.

After obtaining initial estimates of the properties and orientation of the crystal, the beam, the detector, and the goniometer, we split the sequence(s) of images into stills by running laue.sequence to stills, with distinct geometric objects (crystal, goniometer, beam, detector objects) associated with each diffraction image. This step informs downstream programs that the images are stills rather than rotation series data. Although Laue-DIALS currently only supports stills rotation series data (where all data are indexed jointly), we anticipate that this choice will enable the processing of pink serial crystallography data in the near future (where each still represents a random crystal orientation and needs to be indexed independently), with work currently underway to implement the polychromatic indexing algorithm PinkIndexer^24^ natively into DIALS.

### C. Wavelength assignment

To transition from a monochromatic model of the experiment to a polychromatic model, we wrote a new program called laue.optimize indexing, which takes the stills and relaxes the monochromatic constraint to allow for wavelengths anywhere between a user-provided minimum and maximum wavelength. The algorithm iteratively assigns Miller indices and rotates the crystal orientation matrix for the still image to capture as many reflections as possible. Outlier rejection is performed in each iteration by applying the minimum covariance determinant (MCD) algorithm^25^, and the next iteration is performed on the set of inliers calculated in this way. This allows for correcting slight errors in the rotation, but may fail to converge if given large errors such as might be introduced by crystal slippage. Miller indices are assigned by solving a linear sum assignment problem using a cost matrix consisting of the angles between observed and predicted scattering vectors, and wavelengths are determined by the magnitude of the reflection’s scattering vector. The linear sum assignment solver is implemented in SciPy^26^ using the algorithm described by David Crouse.^27^ Rotation updates are estimated by solving the orthogonal Procrustes problem using the algorithm described by Peter Schönemann^28^ also implemented in SciPy^26^. The inferred unit cell of the crystal is kept constant, and is taken from the initial monochromatic estimate. The user may choose to override the initial estimated unit cell with a set of known values, which can correct for errors in the initial estimate resulting from the initial monochromatic lattice search. The output of this program is then a set of DIALS-compatible files with individual wavelengths for each reflection and a set of optimized crystal orientation matrices for each still image.

### D. Geometric Refinement

With the polychromatic indexing solution output by laue.optimize indexing, we can now refine the experimental geometry using the full set of indexed reflections. Laue-DIALS allows for parallel refinement of images. The refinement call, laue.refine, wraps around dials.refine and overrides some parameters to be appropriate for pink data (Table II). In particular, the detector distance and one of the unit cell axes are fixed in order to allow for varying the wavelengths assigned to reflections. If the user wishes to refine either the detector distance or unit cell volume, then the beam wavelengths need to be fixed to converge on a solution. laue.refine then generates a set of DIALS beam objects—one per reflection—all with the same direction, but with individual wavelengths for each reflection. To minimize the memory footprint, these beam objects are generated per image being processed. Once refined, the wavelengths are entered into the reflection data (in the so-called .refl file), and the respective beam objects deleted. Memory limitations can then be handled by reducing the number of parallel processes running, which reduces the number of beam objects instantiated at any particular time. With the parameters supplied, refinement is then run on each still image independently, using the sparse LevMar refinery engine in DIALS^20^ and applying MCD outlier rejection to each still between refinement macrocycles. Spherical reciprocal lattice point models supported in DIALS via the spherical relp model option are used to generate scattering vectors consistent with the vector direction in laue.predict, instead of the default direction which incorporates an Ewald offset. The output files then completely describe an experimental geometry with wavelengths for each observed reflection. Residuals between observed data and predicted reflection centroids are recorded and RMSDs can be visualized using laue.compute rmsds.

### E. Spot prediction

Given a refined experimental geometry, we can now predict the locations of all reflections satisfying the diffraction condition for an image, regardless of whether they were detected during spot finding. To do this, we first generate a set of Miller indices that lie within the envelope bounded by Ewald spheres given by the minimum and maximum wavelengths provided by the user, as well as a maximum resolution (*d*_min_). This set of feasible reflections is then further reduced by filtering harmonics to only the minimal Miller index (for the corresponding reciprocal lattice point closest to the origin of the reciprocal lattice). For the remaining set, wavelengths are then assigned based on the Ewald sphere the index lies on, and then a set of scattered (*S*_1_) vectors is predicted. Those *S*_1_ vectors which point towards the detector are kept.

In order to maximize the accuracy of the predicted reflections, a resolution-dependent bandpass^29^ is then used to filter out improbable reflections based on user-provided tolerance. Probable reflections lie in a region of reciprocal space extending to higher resolution in a spectral intensity-dependent manner as illustrated in Figure 4a. To determine this region, a Gaussian kernel density estimator (KDE) from scipy.stats.gaussian kde is built and trained on the set of resolution and wavelength data corresponding to the observed reflections. To obtain a suitable space for the resolution of each reflection, the resolution is transformed to the square of the distance of each reciprocal lattice point to the origin of the reciprocal lattice (that is, 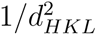), yielding the transformed space shown in panel Figure 4b and c. Harmonic reflections are removed from the training data for the KDE. Treating the resulting smoothed histogram as a probability density function, the full set of feasible reflections are then assigned probabilities based on their resolution and wavelength, with those having a probability lower than the user-provided threshold being removed from the dataset—any observation outside of a probability contour in Figure 4b. The program then outputs a file containing the reflection data for both strong and weak reflections on the image, with observed spots being marked as ”strong” in the reflection table.

**FIG. 4.**
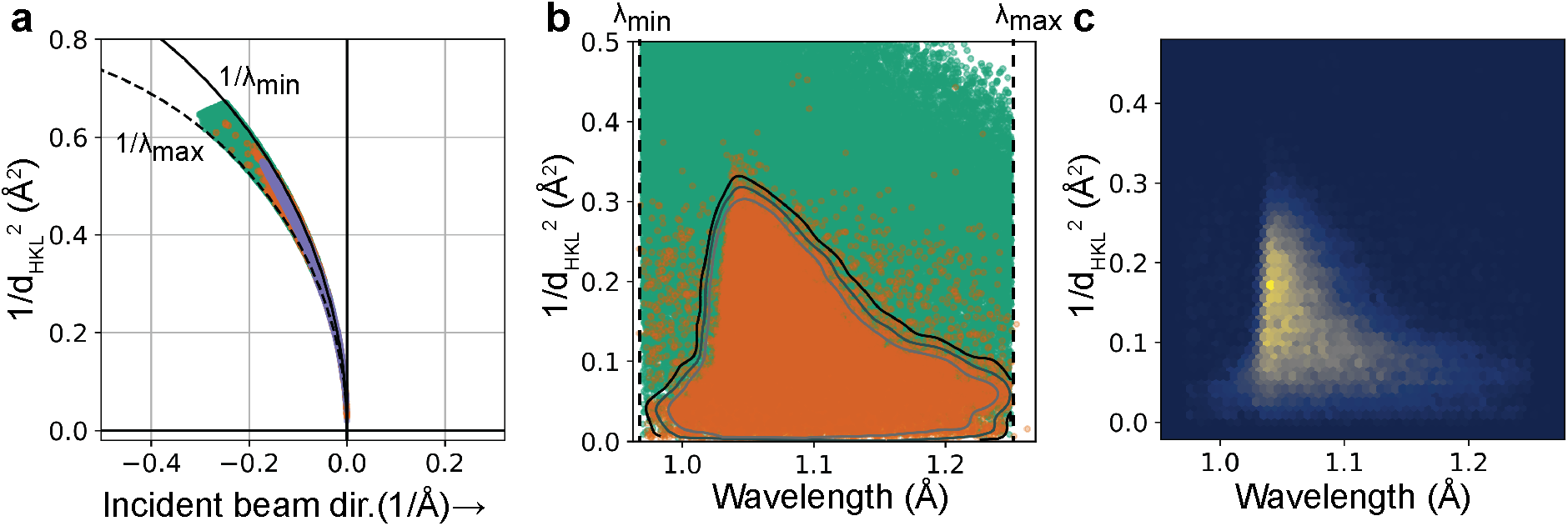
Spot prediction and the use of a resolution-dependent bandwidth. **A.** Ewald diagram for an experimental DHFR dataset, showing the limiting Ewald spheres at the minimum and maximum wavelength (*λ*_min_ and *λ*_max_) used for spot prediction (solid and dashed circular segments, respectively), with predicted reflections (green, random subset of 100,000 reflections), observed reflections (orange), and retained predicted reflections (blue-gray) mapped onto the diagram. **B.** Same predicted and observed reflections in the data representation used for kernel density estimation (KDE). Contours: curves of constant KDE probability density. The retained reflections in panel (a) correspond to the outer contour. **C.** Two-dimensional histogram of the observed (strong) reflections.

### F. Integration

To determine integrated intensities, we use laue.integrate, which implements a variable elliptical integrator similar to the approach described by Ren and Moffat^29^: For each strong reflection, an elliptical profile is built which includes a foreground and background mask for the reflection. Weak reflections then have their profiles built by using a *k*-nearest neighbor approach, averaging the profiles of nearby strong reflections. In the case of overlapping reflections, the integrator assigns all pixels to the nearest reflection centroid for profile estimation and fitting, which avoids integrating the same pixel multiple times. A radius argument can be overridden to only integrate pixels within a given radius of the reflection centroid. The intensities for each reflection are then determined using a summation routine applied to each profile to obtain a set of integrated intensities. The output MTZ file then contains the image numbers, Miller indices, reflection centroids, wavelengths, integrated intensities (both background and foreground), and estimates of the uncertainty of each reflection intensity based on propagating the counting error from each pixel used in summation.

### G. Scaling and merging

The integrated intensities obtained from Laue-DIALS still need to be corrected for variations in the wavelengths giving rise to each reflection (because incident flux, absorption, and scattering depend on wavelength), for the overlap of harmonics, and for other factors not specific to Laue diffraction, such as radiation damage, beam polarization, and variations in diffracting volume during rotation.^30^ Here, we infer and apply the relevant correction factors (or scale factors) using the program Careless, which performs simultaneous inference of the scales of reflections and their merged structure factor amplitudes^13^.

## III. RESULTS

We illustrate the current capabilities of Laue-DIALS using three examples. Detailed implementations of these examples are provided on the Laue-DIALS GitHub page (https://rs-station.github.io/laue-dials/) in the form of didactic Jupyter notebooks accompanying the online documentation. An archive of these tutorials for version 0.4 of Laue-DIALS is available in the accompanying Zenodo deposition (see Data Availability statement). Typical run times are shown in Table III.

**TABLE III.** Processing parameters and time for each dataset. Wall clock time for analysis of three different data sets is provided as a benchmark for performance. All run times are presented in HH:MM:SS format (omitting zero values). For the multi-pass PDZ2 data, Laue-DIALS runtimes are presented for a single pass of 45 images, where 4 CPUs were used for spot finding and indexing, 8 CPUs were used for refinement and spot prediction, and 1 CPU was used for integration. Careless runtime includes all 1064 images. For simulated DHFR data (reference data only) 4 CPUs were used for all processes except for laue.integrate (2 CPUs) and Careless merging (1 GPU). Careless merging times are for training only, and not for running on half-dataset repeats. For the Anomalous HEWL data, 8 CPUs were used for indexing through refinement, 48 CPUs were used for spot prediction, and 64 CPUs were used for integration.

| Dataset | Number of Images | Number of CPUs | Indexing through refinement | Spot prediction | Integration | Careless Merging |
| --- | --- | --- | --- | --- | --- | --- |
| Simulated DHFR | 180 | 4/2 | 12:52 | 8:10 | 1:06:67 | 39:00 |
| Anomalous HEWL | 3,049 | 8/48/64 | 3:07:11 | 6:18:56 | 1:15:37 | 1:21:53 |
| PDZ2 | 1,064 | 4/8/1 | 2:35 | 0:17 | 15:23 | 20:00 |

### A. Processing of simulated ground-truth data

To investigate the accuracy of Laue-DIALS, we created several simulated datasets of diffraction from a crystal of dihydrofolate reductase (DHFR), starting from a deposited dataset (PDB ID 7LVC). The simulations were performed using the program nanoBragg^31,32,33^ as part of the Computational Crystallography Toolbox^34^. With nanoBragg, forward diffraction (the pixel values, given the structure factors and experimental parameters) was calculated according to the classic kinematical theory.^35^ To investigate the sensitivity of Laue-DIALS with respect to various experimental parameters, we simulated 11 pseudo-rotation scans, for each of which we modeled a unique combination of beam divergence, spectral bandwidth, and crystal mosaicity (see Table IV). Each pseudo-rotation scan comprised 180 still exposures, and the crystal was rotated by 1 degree about a fixed axis between exposures, for a total of 180 degrees per scan. Synthetic measurements were recorded in a Rayonix camera format and included realistic background and readout error. Close-ups of representative simulated diffraction images are shown in Figure 2.

**TABLE IV.** Completeness of merged data on simulated data sets. Each simulated data set is analyzed in Laue-DIALS and merged in Careless. Then careless.completeness is run on the merged data to obtain an overall completeness for the data set. Crystal mosaicity, beam divergence, and beam spectral bandwidth are given for each simulated data set. The first row is a reference condition, and the remaining data sets are designed such that they vary from the reference data set by only one parameter. The BioCARS spectrum has a FWHM of approximately 5.5% and is asymmetric.^56^ The other spectra are Gaussians centered on 1.05 Å with the noted bandwidth.

| Mosaicity ($^{\circ}$) | Divergence ($^{\circ}$) | Bandwidth (FWHM %) | Completeness (%) |
| --- | --- | --- | --- |
| 0.1 | 0.1 | BioCARS | 98.5 |
| 0.05 | 0.1 | BioCARS | 99.1 |
| 0.25 | 0.1 | BioCARS | 98.3 |
| 0.5 | 0.1 | BioCARS | 83.1 |
| 0.1 | 0.0001 | BioCARS | 99.0 |
| 0.1 | 0.001 | BioCARS | 99.4 |
| 0.1 | 0.01 | BioCARS | 99.0 |
| 0.1 | 1 | BioCARS | 98.4 |
| 0.1 | 0.1 | 0.5 | 66.8 |
| 0.1 | 0.1 | 1 | 81.6 |
| 0.1 | 0.1 | 2.2 | 96.3 |
| 0.1 | 0.1 | 5 | 98.2 |

The simulated data were processed through the Laue-DIALS pipeline illustrated in Figure 3 and integrated intensities were then merged in Careless (see Methods). We first assessed indexing accuracy. As expected, many more reflections are indexed after polychromatic indexing (laue.optimize indexing) and polychromatic geometry refinement (using laue.refine) than after initial monochromatic indexing, with a concomitant increase in accuracy (Table V). We find that after monochromatic refinement, off-by-one errors on Miller indices are the dominant errors. Polychromatic indexing largely eliminates such errors (compare rows 1 and 2 of Table V). After polychromatic geometry refinement, the primary source of misindexing are harmonic reflections. Of the remaining 163 misindexed reflections 125 (75%) are assigned to harmonics of the correct Miller index. Since Careless considers all harmonics of the assigned Miller index compatible with the wavelength spectrum, such errors will be corrected for in downstream processing. The causes of misindexing of the remaining 0.21% of reflections remain under investigation.

**TABLE V.** Indexing accuracy at key analysis pipeline points. Laue-DIALS was run on simulated DHFR data with known Miller indices. These reflection tables are output by laue.index, laue.optimize indexing, and laue.refine respectively. Note that, (1) polychromatic refinement applies an outlier rejection step, but (2) ultimately all spots are repredicted based on the geometry inferred during polychromatic refinement.

| File | Total Reflections | Correctly Indexed Reflections |
| --- | --- | --- |
| monochromatic.refl | 10,958 | 10,505 (95.87%) |
| optimized.refl | 21,282 | 21,088 (99.09%) |
| poly_refined.refl | 17,722 | 17,559 (99.08%) |

Since the beam is polychromatic, there remains an overall error in the inferred unit cell dimensions. As a consequence, assigned wavelengths can differ from true wavelengths by a multiplicative error, showing in Figure 5 as a systematic deviation from the diagonal. Once accounted for using a scalar correction (blue lines), the assigned wavelengths of 99.91% of reflections in the reference data set coincide with the true wavelengths with an error of less than 0.001 Å. Significant rates of failure only occur for highly mosaic crystals (Figure 5), where outlier reflections’ wavelength errors are likely due to misindexing of harmonic reflections. Consistent with these observations, we find that crystal mosaicity is the main factor impacting merging statistics (Figure 6). Mosaicities higher than 0.1 degrees are uncommon at room temperature but can be induced by freezing of laser- or electric field perturbations or sample handling.

**FIG. 5.**
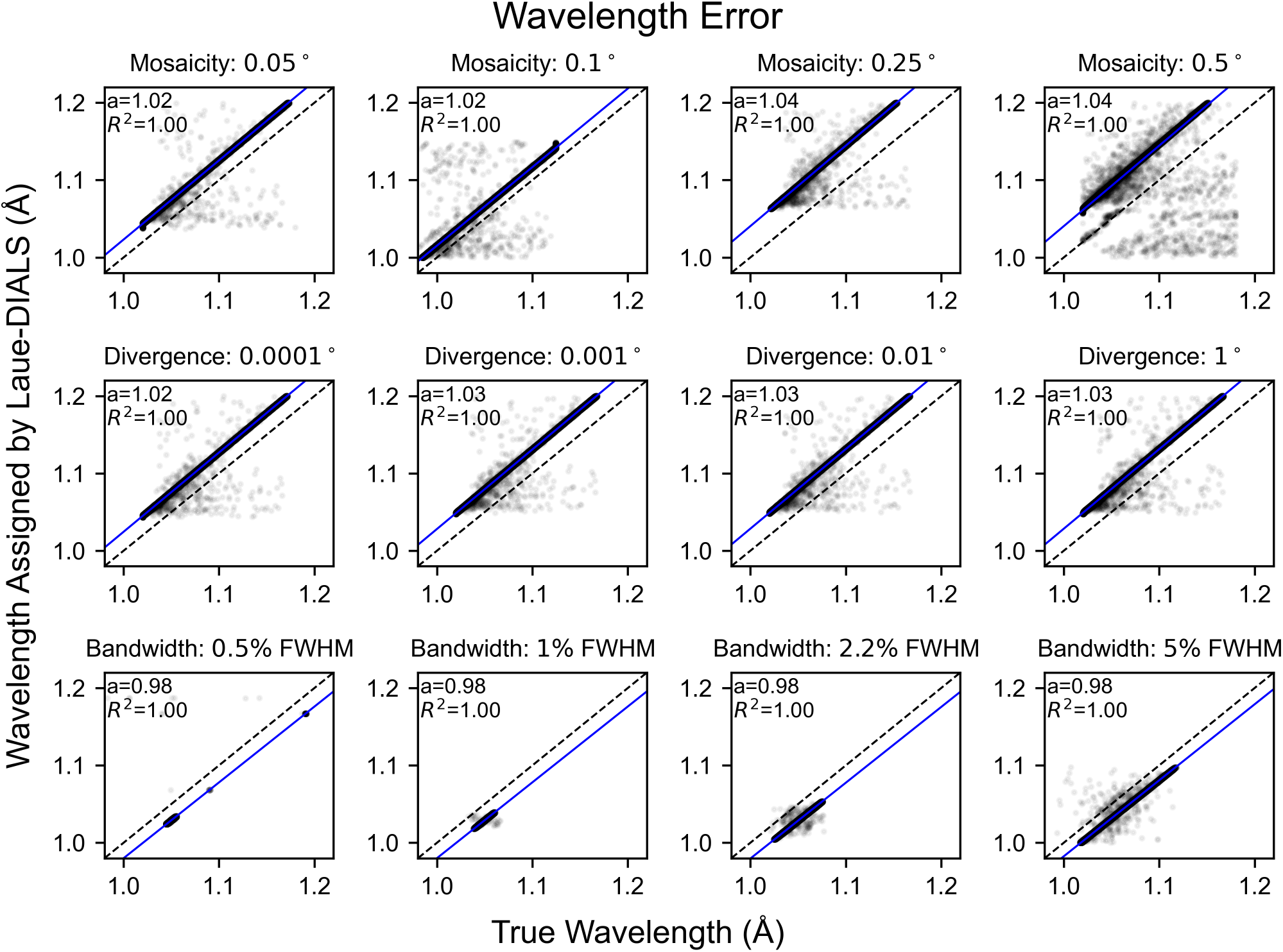
Wavelength errors for processing simulated data. Scatter plots of the simulated wavelengths and wavelengths assigned to reflections by Laue-DIALS. Each subplot has the line y=x plotted in black for reference and includes a least-squares linear regression (blue) with the slope (*a*) and correlation coefficient (*R*^2^). The intercept is set to *b* = 0. Titles for each subplot denote the key parameter difference from the reference data set, which is marked with “Mosaicity: 0.1*^◦^*.” Plots with differing bandwidths use a Gaussian spectrum with the labeled bandwidth. All other plots use a spectrum derived from the BioCARS beamline with approximately 5.5% bandwidth at FWHM. Beam divergence is 0.1*^◦^* unless otherwise specified.

**FIG. 6.**
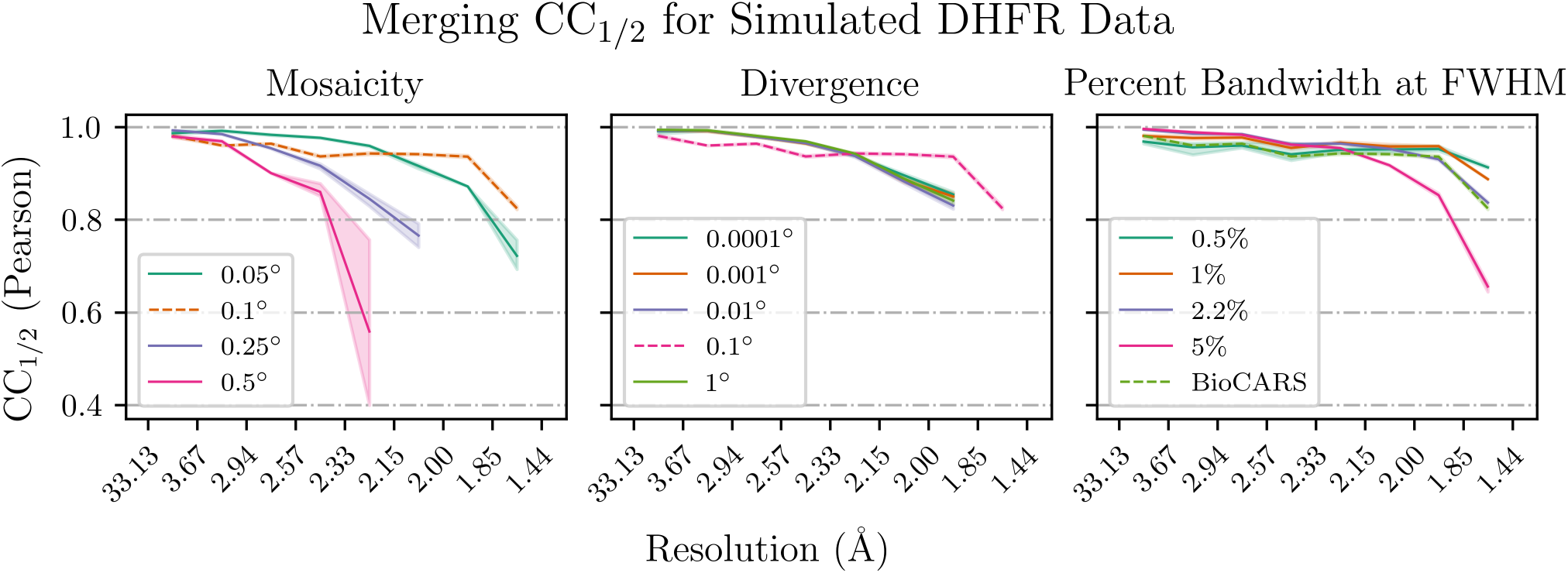
Merging statistics for simulated data. The Pearson *CC*_1_*_/_*_2_ binned by resolution is plotted for each varied parameter in the simulated data. Within each subplot, all parameters except for the labeled parameter are held constant. The dotted line represents the reference data set that is common to all three subplots. Bandwidths labeled as a percentage are Gaussian spectra centered on the BioCARS spectrum with bandwidths as labeled.

Finally, the simulated data make clear the benefit of treating polychromatic data as polychromatic: if we were to only use a monochromatic nominal wavelength for predicting centroids compared to our polychromatic method, large inaccuracies in spot centroid position would result. In Figure 7, we use an adapted form of cctbx.xfel.detector residuals to plot spot centroid residuals within 64 regions (virtual panels) of the detector when predicting spot centroids using either assigned wavelengths (panel a) or using a nominal wavelength (panel b), with coloring indicating the angular error.

**FIG. 7.**
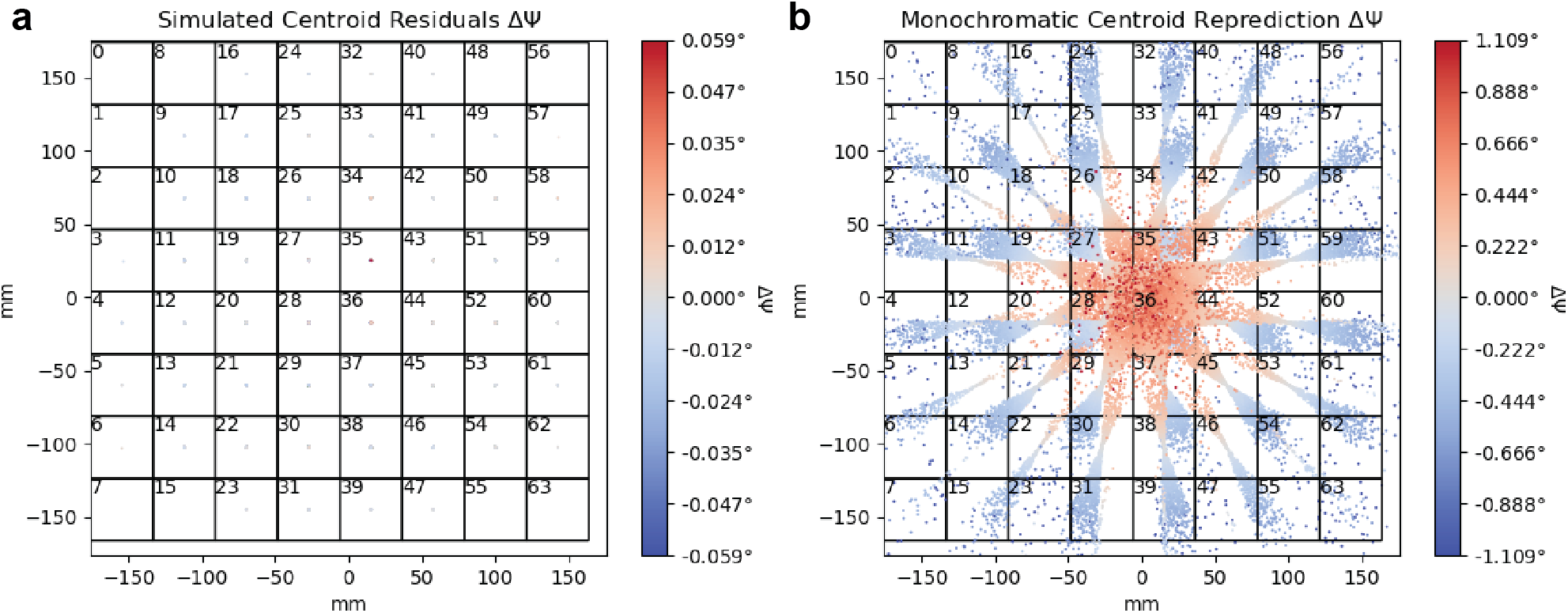
Centroid residuals after geometric refinement for reference simulated data set. **A.** Centroid residuals on the detector binned into 64 equal panels. The colormap denotes the residual error in the refined rotation angle, Δ*ψ*. Wavelengths are relaxed to match spot centroid location, leading to near-zero radial residuals. **B.** Centroid residuals on the detector binned into 64 equal panels. Similar to panel A, but with spot centroids repredicted using the monochromatic nominal beam wavelength prior to calculating residuals. Wavelengths are not relaxed here, leading to high radial residuals illustrating the difficulties of applying monochromatic indexing routines to these data. Plots were produced using a modified version of cctbx.xfel.detector residuals.

### B. Analysis of anomalous signal

Anomalous diffraction is a vital component of many macromolecular crystallographic experiments. In addition to enabling experimental phasing^36,37^, anomalous signal can be used to distinguish biologically relevant ions^38^. Anomalous peak heights can also serve as a sensitive reporter of data processing accuracy^39^. Such anomalous scattering may come either from atoms natively present in the protein, such as sulfur, or from atoms or ions soaked into the crystal. To examine whether Laue-DIALS could resolve anomalous signal from single-crystal polychromatic data, we collected a pseudo-rotation series of 3049 frames on a HEWL crystal soaked with sodium iodide. The data were collected at ambient temperature (about 295 K) with the standard BioCARS X-ray beam (1.02-1.20 Å, with 1.05 Å peak intensity wavelength). Notably, this peak wavelength is far from the X-ray absorption edges of either iodine or sulfur. We processed the data with Laue-DIALS (see Methods and the accompanying Jupyter notebook) before scaling with Careless^13^ and refining with Phenix^40^.

The resulting anomalous map showed clear peaks on all 10 native sulfur atoms as well as 5 ordered iodide ions (Figure 8a; confirmed by comparison to a monochromatic reference dataset, see Methods). The crystal diffracted well, with a CC_1/2_ value of over 0.975 at 1.7 Å (Figure 8b); the CC_anom_ value remained above the 0.3 throughout that same range (Figure 8c). Since this is a relatively large single-crystal dataset, we also scaled and merged subsets of the data in Careless to examine the strength of the anomalous signal from subsets of the data. Whether the 15 anomalous scattering atoms were considered individually (Figure 8d) or averaged by element type (Figure 8e), the results remained consistent. While the anomalous peak height increases with increasing amount of diffraction data, nearly all of the signal could be obtained in under 1500 frames, and based on the provided fits, the anomalous signal of S and I atoms reaches 50% of its asymptotic value within 180 frames (about 85 frames for I and 156 frames for S atoms).

**FIG. 8.**
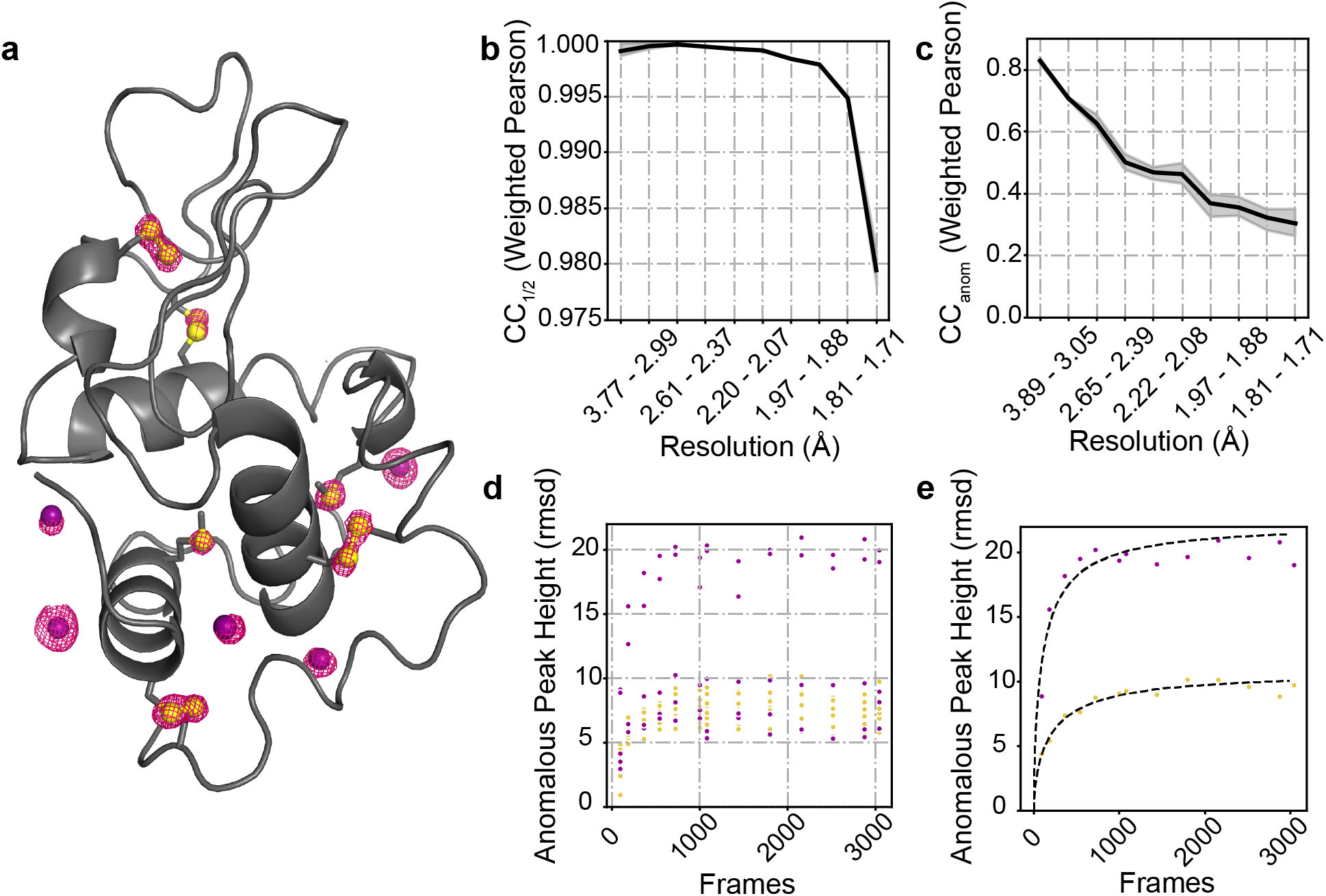
Sulfur and Iodine Anomalous signal from a Hen Egg White Lysozyme (HEWL) crystal soaked with NaI. **A.** Anomalous peaks at 1440 frames contoured at 4 *σ* (within 1.6 Åof model atoms). Phases and model: PDB ID 9B7C. Yellow spheres: sulfur atoms; purple spheres: iodide ions. **B.** *CC*_anom_ as a function of resolution bin after 1440 frames. **C.** *CC*_half_ as a function of resolution bin after 1440 frames. **D.** Anomalous peak heights for each of the five ordered iodine atoms (purple) and ten sulfur atoms (yellow) present in the HEWL structure. **E.** Anomalous peak heights vs frame numbers for the IOD 4 (purple) and Cys80 (yellow) atoms. Fits were obtained as described in ref.^53^ and take the form 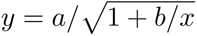, where *a* and *b* are fitting parameters and *x* the number of frames.

### C. Analysis of time-resolved diffraction data

Conventional fixed-target, time-resolved Laue diffraction data are typically collected in multiple passes (either on the same or several crystals), with interleaved collection of the unperturbed (“OFF”) data and perturbed (“ON”) data taking place at each angle before moving on to the next angle. These angular steps can be large (4-6°) to ensure even coverage of reciprocal space before a crystal becomes damaged.

Here we illustrate the processing of such time-resolved diffraction data with Laue-DIALS for data from an EF-X experiment (electric-field-stimulated time-resolved x-ray crystallog-raphy). This EF-X dataset of the second PDZ domain of human LNX2^41^ contains sixteen image series, each obtained from a phi angle scan. Each pass has four electric-field timepoints—off, 50ns, 100ns, and 200ns of electric field. The workflow is illustrated in Figure 9a and described in detail in an accompanying Jupyter notebook. This workflow ensures consistent geometry in all processing steps.

**FIG. 9.**
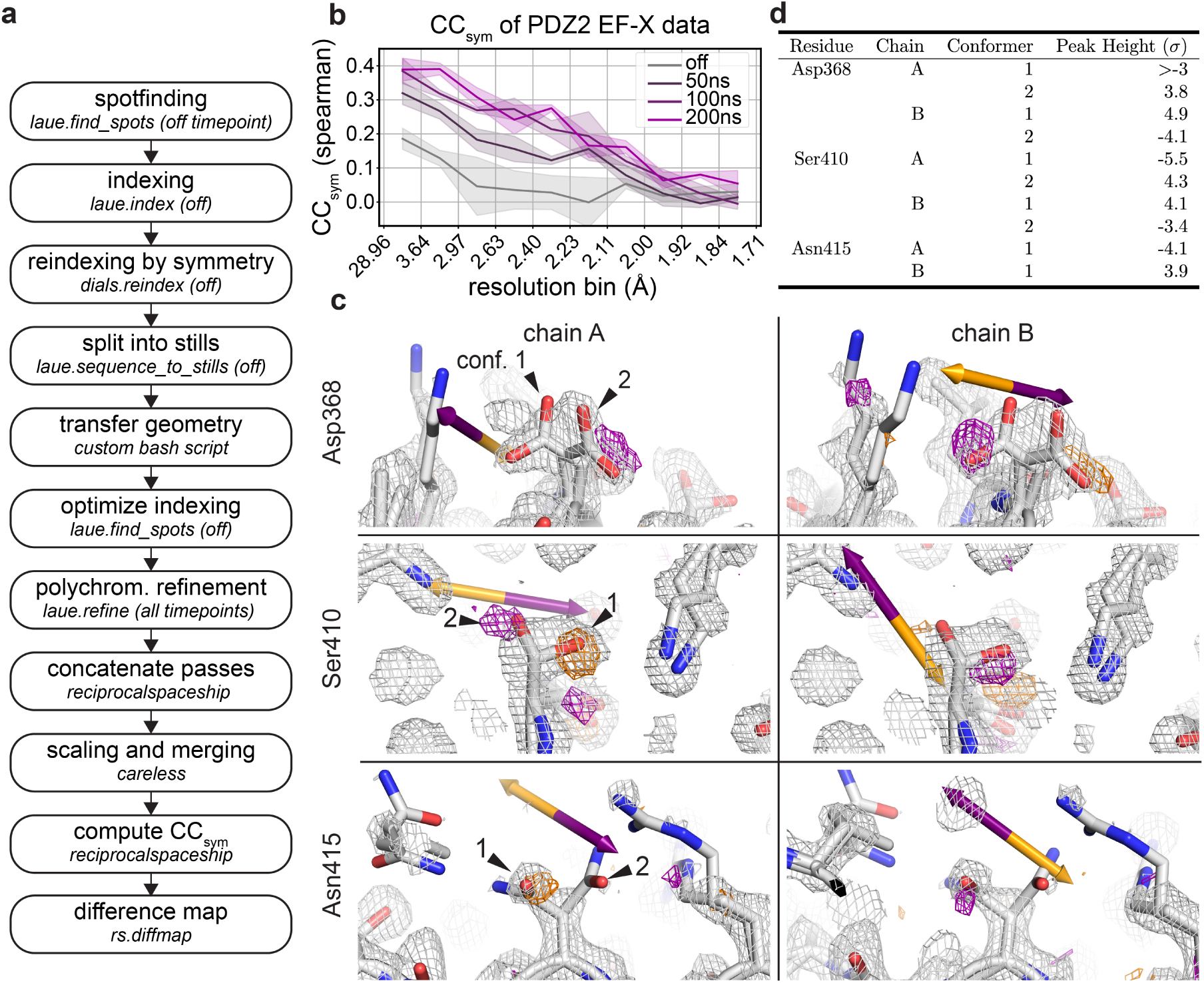
Time-resolved signal in an EF-X dataset processed with Laue-DIALS. The dataset is from an EF-X experiment on PDZ2 with one OFF and three ON timepoints, originally reported in Hekstra et al. 2016. **A.** Flowchart of the processing workflow. **B.** Plot of *CC*_sym_ as a function of resolution bin for each timepoint. **C.** Weighted ON*−*OFF isomorphous difference maps showing electric-field induced sidechain motions. Orange arrows depict the direction of the electric field, and purple arrows depict the opposite direction. Orange density represents a decrease in density in the ON map compared to the OFF map, and purple density represents increased density in the ON map. Maps are contoured at 3*σ* and carved within 1.5 Åof shown atoms. There is an increase in electron density on Asp368 carboxylate conformer 2 in one symmetry mate (chain A, top left) and conformer 1 in the other (chain B, top right). There is an increase in electron density on Ser410 hydroxyl conformer 2 in one symmetry mate (chain A, middle left) and conformer 1 in the other (chain B, middle right). There is a decrease in electron density on Asn415 carboxamide conformer 1 in one symmetry mate (chain A, bottom left) and increase in the other (chain B, bottom right). **D.** Heights of difference map peaks in (C).

As an overall measure of time-resolved signal in reciprocal space, we use *CC*_sym_, which quantifies symmetry breaking due to the electric field, analogous to how *CC*_anom_ quantifies deviations from symmetry imposed by Friedel’s law due to anomalous signal by measuring the correlation of structure factor amplitude differences between Friedel mates estimated from separate halves of the data^42^ (see Methods). *CC*_sym_ clearly indicates the presence of a signal increasing with the duration of the electric field, consistent with previous observations (Figure 9, ref.^41^).

Weighted difference maps highlight electric-field dependent motions (Figure 9) consistent with those observed previously^41^ and highlighted here for residues Asp368 and Ser410: for Asp368 we observe a shift in rotamer equilibrium with side chain motion against the applied electric field, consistent with naive expectation for the motion of a negatively charged group in an electric field (indicated by an orange arrow; Figure 9, top left and right). Additionally, Ser410 in both chain A and chain B moves against the electric field (Figure 9, middle), while Asn415 in both chain A and chain B moves with the electric field (Figure 9, bottom). Both of these observations are as expected based on previous observations.^41^ Of the difference map peaks highlighted in Figure 9, peak heights are above 3*σ* in Coot (Figure 9)^43^. We conclude that time-resolved signal can be recovered by analysis with Laue-DIALS.

## IV. DISCUSSION AND CONCLUSIONS

We have described a computational framework, Laue-DIALS, for open-source processing of polychromatic X-ray diffraction patterns, primarily intended for processing macromolecular data. Laue-DIALS can be installed as an add-on package for DIALS, a general framework for processing diffraction data, and builds on its code base. Like DIALS, Laue-DIALS is open-source and free, and welcomes contributions and community involvement. Laue-DIALS also inherits DIALS’ modular architecture such that new algorithms can be swapped with relative ease.

This flexibility also allows for a series of planned further improvements and extensions. In particular, the current indexing algorithm was designed for monochromatic data. In future work, we intend to incorporate a natively polychromatic indexing algorithm such as PinkIndexer^24^ or machine learning-based algorithm like LaueNN^44^. Recent work shows the substantial benefit of Laue diffraction for serial crystallography applications^45,46,47,48^. PinkIndexer can index individual frames, potentially removing the primary obstacle to processing serial Laue diffraction data with Laue-DIALS. In the current approach, after indexing and initial inference of the crystal lattice and orientation, the data are already otherwise treated as stills. We further anticipate that natively polychromatic geometric refinement, currently under development in DIALS for processing of polychromatic neutron diffraction data (https://github.com/dials/dials/pull/2662), can improve speed and accuracy of data processing.

## V. METHODS

### A. Simulated Diffraction Data

Simulated diffraction data were generated using nanoBragg.^31,32,33^ For simulations, we assumed a parallelepiped crystal domain consisting of 100 unit cells along each unit cell vector. The cubed root of the resulting crystal domain volume was 5.4 microns (using the unit cell taken from PDB 7LVC), and this domain size defined the characteristic profile of each Bragg reflection. However, to simulate scattering from a crystal with 100 micron thickness, we amplified the scattering by a factor of 3.6 *×* 10^5^. The incident beam spot size was set to 10 microns, and the total photons per exposure was 5 *×* 10^11^. In addition to the crystal diffraction, we modeled scattering through 2.5 mm of water and 5 mm of air. We simulated data onto a detector of area 340 mm x 340 mm, with 3840 pixels along each dimension, making a pixel size of 0.088 mm. The crystal-to-detector distance was 200 mm. To simulate energy dispersion, a photon energy spectrum with 5 eV resolution (10 eV for the Gaussian spectra as they were smoothly varying) was created spanning the energy range, leading to 322 discretely sampled energies per exposure. For each energy, we modeled 12 beam vectors spanning a cone of divergence. And for each exposure, to simulate angular mosaic spread, we perturbed the nominal crystal orientation 100 times (according to the desired mosaic spread), and averaged the diffraction from each perturbed crystal. These discrete calculations resulted in 322 x 12 x 100 = 386,400 simulation steps per exposure, warranting GPU acceleration. Diffraction was simulated into each square pixel. Pixel gain was set to 0.7 ADUs (area detector units) per photon. In addition to Poisson noise on the number of photons arriving at each pixel, gain calibration noise (a randomly sampled multiplicative factor on the per pixel gain) and readout noise (a per pixel, per exposure randomly sampled additive factor on the amplified signal) was modeled. Per-pixel gain calibration terms were drawn from a normal distribution centered on 1, with a standard deviation of 0.03. Per-pixel, per-exposure readout noise terms were drawn from a normal distribution centered on 0, with a standard deviation of 3 ADUs. Finally detector point spread was modeled using values typical for Rayonix cameras.^49^

For each simulated data set of 180 images, the image data were imported using a custom Python script that built DIALS objects according to the simulated parameters and read the synthetic image data from the CBF files. These data were then processed through the standard pipeline (Figure 3). The same set of parameters is used in each analysis, with wavelength limits of 0.9 and 1.2 Å used for all analyses. The known unit cell for the simulated DHFR crystal was input into laue.index and allowed to undergo scan-varying refinement before being split into stills. Based on analyses of experimental DHFR data, a resolution cutoff of 1.4 Å was supplied to laue.optimize indexing and laue.predict. Each image was then integrated using the variable elliptical algorithm implemented in laue.integrate and combined into MTZ files containing the integrated intensities for each separate simulated data set. These data sets were then merged with Careless^13^, and supplying the wavelengths output from laue.refine for each reflection.

We attached fractional Miller index data to pixels of the reference simulated data set, and used these to determine misindexing rates throughout the pipeline. We analyzed the output files of laue.index, laue.optimize indexing, and laue.refine by locating the pixel of the spot centroid for each indexed reflection, rounding the fractional Miller index in the associated image data, and checking for consistency with the assigned Miller index in the reflection table. Wavelength errors for each simulated data set were similarly calculated by comparing reflection wavelengths to the intensity-weighted average wavelength associated with the pixel of the spot centroid.

To compare monochromatic and polychromatic analyses of these data, we processed the data through geometric refinement as in Section III A and applied an additional set of analyses to the output of laue.refine. Firstly we averaged the respective detector objects for each image into a single detector object for the data set. A custom Python script then split these detectors into sets of 8×8 panels. The resulting data were passed to a custom variant of cctbx.xfel.detector residuals twice – once with repredict input reflections=False and once with repredict input reflections=True. The former case plotted detector residuals in each panel using the wavelength estimates given in Laue-DIALS, while the latter case repredicted spot centroids using the single nominal wavelength provided to laue.index.

### B. Collection and Analysis of Lysozyme Anomalous Data

Crystals of hen egg white lysozyme (HEWL) were grown and soaked with sodium iodide as described in ref.^50^. We collected a 3049-frame Laue dataset at BioCARS (Advanced Photon Source (APS) beamline 14-ID-B) on a single HEWL crystal soaked with sodium iodide. Data were collected at room temperature (about 295 K). On the same day, we collected a monochromatic 295 K data set on an NaI-soaked crystal from the same batch at 1.0375 Å at beamline 24-ID-C at NE-CAT, APS. Both datasets will be described in further detail in a future manuscript.

A complete Laue-DIALS data analysis protocol is described in an accompanying Jupyter notebook (see Data Availability statement). Briefly, we processed the full dataset in Laue-DIALS, yielding an integrated MTZ file. This MTZ file was split into two subsequent MTZ files containing either the positive or the negative Friedel mates using reciprocalspaceship^51^. Both MTZ files were scaled with Careless using a bivariate prior on structure factor amplitudes^13^ (see also https://github.com/rs-station/careless-examples/). The Friedel mates for each dataset were then recombined after scaling and merging. Phases for determination of anomalous peak heights were obtained from a model refined against the monochromatic reference dataset. This model was built by experimental phasing using AutoSol^52^, AutoBuild, and phenix.refine^40^, and has been deposited as PDB ID 9B7C. To assess the effect of data redundancy on anomalous signal, we repeated this analysis starting from integrated intensities from the first 90, 180, 360, 540, 720, 999, 1080, 1440, 1800, 2160, 2520, 2880, or 3049 (complete) diffraction frames.

### C. Analysis of time-resolved data

A complete data analysis protocol is described in an accompanying Jupyter notebook (see Data Availability statement). Briefly, these single-crystal, fixed-target data were collected in an interleaved fashion, collecting OFF (no voltage applied) and ON (X-ray exposure at multiple delays from the start of a high-voltage pulse) at each crystal rotation angle, followed by sample rotation. We first process the four consecutive OFF passes independently by running the full Laue-DIALS pipeline through integration. After laue.index, we check for consistency of the inferred geometry and, if necessary, apply the (*−x, y, −z*) symmetry operation of the C2 space group using dials.reindex. This geometry is then used for laue.sequence to stills. At this point, we transfer the OFF geometry to the corresponding ON image series and proceed with geometry optimization, polychromatic refinement, and integration for the ON data. Using the included custom scripts based on reciprocalspaceship^51^, we combine passes for each timepoint and prepare MTZ files in both the original and reduced-symmetry space groups (see Hekstra et al.^41^). These .mtz files are reduced together using Careless.^13^

*CC*_sym_ was introduced by Greisman et al.^42^ and calculated using a custom script adapted from rs-booster (https://rs-station.github.io/), an add-on package to reciprocalspaceship. As expected, there is a higher resolution-dependent *CC*_sym_ for later timepoints. Weighted difference maps between the 200ns and off timepoints were calculated using tools from reciprocalspaceship and rs-booster.

## ACKNOWLEDGMENTS

We thank Drs. Rama Ranganathan and Keith Moffat for advice and encouragement, Dr. David Waterman for stimulating discussions on polychromatic refinement, Michael Socolich (University of Chicago) for providing the NaI-soaked lysozyme crystals, and to Robert Henning and Insik Kim for remote collection of diffraction data for the HEWL data set. This work was supported by a research agreement between BioCARS and the University of Chicago, a Burroughs-Wellcome Fund CASI award (KMD); a National Science Foundation Graduate Research Fellowship (to HKW, DGE 2140743), an NIH grant (R35-GM151988; NKS), a DOE ICDI grant (DE-SC0022215; ASB), and the NIH Director’s New Innovator Award (DP2-GM141000; DRH). Finally, we gratefully acknowledge past and present efforts of the community to make the routine, accurate processing of Laue diffraction data a reality. KMD is currently employed by the Linac Coherent Light Source (LCLS) at the SLAC National Accelerator Laboratory, an Office of Science User Facility operated for the U.S. Department of Energy Office of Science by Stanford University. KMD is supported by the U.S. Department of Energy, Office of Science, Basic Energy Sciences under Contract No. DEAC02-76SF00515.

## VI. AUTHOR DECLARATIONS

### B. Author Contributions

**Rick A. Hewitt:** Data Curation (equal); formal analysis (equal); software (lead); validation (equal); visualization (equal); writing - original draft (equal); writing - review and editing (equal). **Kevin M. Dalton:** Conceptualization (lead); data curation (equal); formal analysis (lead); investigation (equal); methodology (lead); software (equal); supervision (equal); visualization (supporting); writing - original draft preparation (equal); writing - review and editing (equal). **Derek Mendez:** Data curation (equal); formal analysis (equal); investigation (equal); software (supporting); visualization (equal); writing - review and editing (equal). **Harrison K. Wang:** Data curation (equal); formal analysis (equal); investigation (equal); software (supporting); validation (equal); visualization (equal); writing - original draft preparation (equal); writing - review and editing (equal). **Margaret A. Klureza:** Data curation (equal); formal analysis (equal); investigation (equal); software (supporting); validation (equal); visualization (equal); writing - original draft preparation (equal); writing - review and editing (equal). **Dennis E. Brookner:** Data curation (equal); formal analysis (supporting); investigation (equal); software (supporting); validation (equal); visualization (supporting). **Jack Greisman:** Data curation (equal); investigation (equal); software (supporting); writing - review and editing (equal). **David McDonagh:** Formal analysis (supporting); validation (equal); visualization (supporting); writing - review and editing (equal). **Vukica S^̌^rajer:** Conceptualization (equal); data curation (equal); investigation (equal); validation (equal); writing - review and editing (equal). **Nicholas K. Sauter:** Conceptualization (equal); writing - review and editing (equal). **Aaron S. Brewster:** Conceptualization (equal); data curation (supporting); formal analysis (equal); methodology (equal); software (supporting); validation (supporting); visualization (supporting); writing - original draft preparation (equal); writing - review and editing (equal). **Doeke R. Hekstra:** Conceptualization (lead); data curation (equal); formal analysis (equal); funding acquisition (lead); investigation (equal); methodology (equal); project administration (lead); software (supporting); supervision (lead); validation (equal); visualization (equal); writing - original draft preparation (equal); writing - review and editing (equal).

## VII. DATA AVAILABILITY

Full documentation of the Laue-DIALS functionality and worked examples are described at https://rs-station.github.io/laue-dials/. Real data are available on SBGrid for the HEWL (https://data.sbgrid.org/dataset/1118/), PDZ2 (https://data.sbgrid. org/dataset/1116/), and DHFR (https://data.sbgrid.org/dataset/1117/) data sets, with simulated DHFR data available upon request from the corresponding authors. Scripts and notebooks used in this manuscript are deposited in an accompanying Zenodo deposition (https://zenodo.org/records/12761162), which includes a script for downloading data from SBGrid.

